# *C. elegans* small heat-shock protein HSP-12.6 has a highly specialized protective function towards muscle thick filaments *in vivo*

**DOI:** 10.64898/2026.05.17.725775

**Authors:** Abigail Fern, Jasmine Alexander-Floyd, Anhelina Volchok, Susan M. Cahill, Srikar Donepudi, Jana Smuts, Tali Gidalevitz

**Affiliations:** Biology Department, Drexel University, Philadelphia, PA. USA; SK Pharmteco, PA, USA; Medical Sciences Division, University of Oxford, Oxford, UK; Department of Medicine, Rutgers New Jersey Medical School, Newark, NJ, USA; Neurobiology and Anatomy Department, Drexel University, Philadelphia, PA, USA

**Keywords:** small heat shock proteins, HSP-12.6, muscle proteostasis, myofilaments, chaperones, protein aggregation

## Abstract

Small heat-shock proteins (sHSPs) are an ancient and diverse class of molecular chaperones, acting as a first line of defense against proteotoxic stresses. While the canonical sHSPs prevent uncontrollable aggregation of a broad range of non-native substrates, a subset of sHSPs do not exhibit this broad activity *in vitro*, and their functions *in vivo* are poorly understood. Interestingly, several such sHSPs are selectively expressed in muscle tissues, including by myogenic programs, indicating likely functional roles. We examined *in vivo* function of *C. elegans* HSP-12.6, which possesses no chaperone activity *in vitro* but regulates lifespan, and is developmentally induced in the muscles of long-lived dauer animals. We found that HSP-12.6 exhibits exceptional selectivity in protecting the muscle function against folding or assembly mutations in thick filament proteins, but not in thin filament or non-filament proteins. This reflected its exclusive chaperone-like binding to the healthy myosin-containing thick filaments, and to their aggregates. HSP-12.6 did not bind other muscle structures or aggregates, including those of thin filaments, and retained its selectivity to either healthy thick filaments or their aggregates when challenged with a toxic aggregation-prone polyQ protein. Our data establish HSP-12.6 as a highly-selective myoprotective chaperone, with client spectrum distinct from other sHSPs.

## Introduction

Small heat-shock proteins (sHSPs) are ATP-independent molecular chaperones ^1,2^ present in all kingdoms of life ^3^. They are defined by a highly conserved α-crystallin domain, flanked by divergent and largely unstructured N- and C-terminal extensions ^4,5^. sHSPs preserve cellular functions and increase survival under proteotoxic stress conditions, and are required for both the normal lifespan and its extension in animal models (*e*.*g*. in insulin/IGF signaling mutants) ^6,7^. Their defects are linked to many human diseases, including neuropathies and neurodegeneration, skeletal and cardiac muscle diseases, cataracts, and cancer ^8^. Two types of sHSPs are distinguished in metazoans: those with broad substrate specificity, referred to as generalists, and those with restricted substrates (or clients), known as specialists ^5,9^. Most of our substantial knowledge on the mechanisms and functions of sHSPs comes from the generalists, including the archetypal sHSPs - HSPB5/αB-crystallin and HSPB1/Hsp27. *In vitro*, they typically equilibrate between dimers or tetramers and large polydispersed oligomers, depending on their environment, and prevent aggregation of unfolded model substrates ^4,5,10^. *In vivo*, their expression is controlled by stress, and they function by recognizing a variety of non-native and destabilized proteins, either sequestering them into large aggregates or forming smaller soluble complexes ^4,5,10^. These functions shield the cellular proteome from inappropriate interactions during stress, and allow non-native proteins to be refolded by ATP-dependent chaperones, or targeted for degradation, once the stress is resolved ^11-14^. The generalist sHSPs can also function outside of the global stress paradigm, for example by regulating intermediate filaments ^15^.

In contrast, the functions and mechanisms of the specialist sHSPs are less clear. First, their expression is often independent of stress, and they are instead expressed in tissue-specific or developmental stage-dependent patterns. Interestingly, seven of the ten human sHSPs are either selectively expressed or enriched in the muscle and cardiac tissues, including three that are directly induced by myogenic transcription factors; this is also true in the model organisms, suggesting that muscles may have specific requirements for sHSPs ^16-19^. Second, few examples of their native physiological clients are known, including control of dimerization of the actin-interacting protein Filamin C by HSPB7/cvHSP (cardio-vascular HSP) in striated muscles ^20^, and regulation of protein interactions of 14-3-3 dimer by HSPB6/Hsp20, to control relaxation of smooth muscles ^21,22^.

Strikingly, specialist sHSPs often show weak or no activity towards unfolded substrates. For example, an *in vitro* study that compared all ten human sHSPs (HSPB1 through HSPB10) found that only the generalists - HSPB1/Hsp27 and two α-crystallins, HSPB4 and HSPB5, showed promiscuous activity, while others were assay- or substrate-selective, and HSPB7 was inactive in all assays ^23^. Similarly, in a cell-based study, only the generalist human sHSPs were active in stabilizing luciferase against thermal unfolding ^24^. Surprisingly, in the same study, HSPB7 was the only sHSP able to suppress aggregation and toxicity of expanded polyglutamine proteins, both in cells and in *Drosophila* model ^24^, suggesting that the apparent lack of activity of the specialist sHSPs in chaperone assays may reflect their selectivity to a narrow set of client proteins.

The puzzling apparent lack of activity is not unique to the human specialist sHSPs – several *in vitro* studies showed that while sHSPs from *C. elegans* HSP-16 family function as promiscuous chaperones, the proteins from HSP-12 family were completely inactive in same assays ^25-27^. A study in yeast supported a generalist function for the HSP-16 family proteins also in cells, by showing that they are able to substitute for the yeast Btn2 and thus function as sequestrases in a heterologous system; in contrast, proteins from the HSP-12 family were inactive as sequestrases ^27^. Furthermore, unlike HSP-16s (and some other sHSPs), none of the HSP-12 family proteins are enriched in aging-induced aggregates ^28,29^. Yet, genetic experiments in *C. elegans* unambiguously show that HSP-12 family proteins have important *in vivo* functions ^7,19,30,31^. Thus, understanding the *in vivo* chaperone functions of the specialist sHSPs will require uncovering their physiological clients.

Here we use an HSP-12 family member, HSP-12.6, to test whether it can function as a molecular chaperone *in vivo. hsp-12*.*6* is highly upregulated in the long-lived and exceptionally stress-resistant developmental stage called dauer ^32^, and in fact was the most highly expressed dauer-specific transcript by serial analysis of gene expression (SAGE) ^33^. While it is not induced by proteotoxic stresses ^26^, it is regulated by DAF-16 transcription factor, an orthologue of mammalian FOXO3 that functions in the insulin/IGF signaling pathway that controls lifespan ^7,34^, and is also induced by fasting or starvation ^35,36^. In fasted animals, induction of *hsp-12*.*6* is restricted to the muscle tissues and few individual neurons; *hsp-12*.*6* is also strongly expressed in the vulva muscles in well-fed animals ^36^. Deletion or downregulation of *hsp-12*.*6* reverses the lifespan extension of insulin/IGF receptor (*daf-2*) mutants ^7,30^, and decreases osmotic stress resistance of IP3K/*age-1* mutant animals ^31^. These data combined point to a possible protective role in maintenance of proteostasis. However, HSP-12.6 was found inactive in chaperone assays ^26^.

We find that fluorescent fusions of HSP-12.6 localize exclusively to the myosin-containing thick myofilaments in the body-wall muscle cells, both when naturally induced by dauer or fasting, and when ectopically expressed in non-stressed animals. In contrast, HSP-16.48 protein showed enrichment on muscle attachment structures, including M-lines and dense bodies (the worm equivalent of z-disks), as well as diffuse localization, similar to several mammalian sHSPs ^37,38^. The localization of HSP-12.6 to the thick filaments did not depend on the tagging protein, was exclusive of thin filaments, attachments, and other muscle organelles, and we were unable to detect diffuse protein. To understand whether this HSP-12.6 localization reflected its function, we introduced individual mutations that interfere with either folding, assembly, or organization of myofilament proteins, or affect other muscle proteins. We found that the presence of HSP-12.6 improved muscle function and reversed severe phenotypes for all tested folding or assembly thick filament mutations, suggesting a chaperone function. In agreement, HSP-12.6 did not show protection when thick filament proteins were deleted, or when a non-myofilament protein was mutated. Strikingly, HSP-12.6 did not show any protection against mutations in thin filament proteins, and in fact worsened some phenotypes when expressed in animals without thick filament folding/assembly defects. We then asked whether its binding properties agreed with a chaperone function. FRAP analysis showed that it indeed bound thick filaments transiently, with slow recovery, while imaging of folding and assembly mutants showed preferential binding to the aggregates formed by the mutant thick filament proteins, relative to the remaining filaments. Like its protective function, the binding to aggregates was highly selective: HSP-12.6 did not co-localize with aggregates of thin filament proteins. Moreover, when presented with a choice of thick-filament aggregates *vs* another aggregation-prone protein (polyQ) in the same cell, HSP-12.6 exclusively localized to the myofilament aggregates. Our data thus show a strikingly specialized myo-protective function for HSP-12.6 sHSP in *C. elegans* body-wall muscle cells. Its selectivity towards myosin-containing thick filaments is unique among the known sHSP, whose sarcomeric functions are typically directed to thin filaments, their interacting proteins, and their attachments.

## Results

### HSP-12.6 uniquely localizes to the thick filaments in the body wall muscles

GFP-tagged HSP-12.6, driven by a long p*hsp-12*.*6*(5Kb) promoter, was previously established to study its expression, which was detected predominantly in the vulva muscles of well-fed adult animals, and additionally induced in the body-wall muscles (BWMs) in fasted animals ^36^. Because reported *hsp-12*.*6* levels in dauers are very high ^32,33^, we asked if it has a broader expression pattern at this developmental stage. Using the same transgene, we indeed observed very strong expression of HSP-12.6::GFP in dauer animals, but it was still restricted to the muscle cells (Fig. 1A), plus occasional individual neurons. A closer examination showed that HSP-12.6::GFP protein in the body-wall muscle cells (BWMs) was localized in a myofilament-like pattern (Fig. 1A), with no discernable diffuse fluorescence. To better understand this localization, we generated animals carrying integrated low copy-number transgene expressing HSP-12.6::mCherry protein from a muscle-specific promoter (myosin heavy chain A promoter, p*myo-3*), to allow for examination of HSP-12.6 localization in healthy, non-starved muscle cells. We first confirmed that the two proteins behaved similarly in fasted animals. The GFP- and mCherry-tagged proteins showed identical localization to the filaments that were positioned in between the dense bodies (DBs) and overlapping the M-lines (Fig. 1B). Since this is a characteristic position of the myosin-containing thick filaments in striated muscles (see scheme in Sup. Fig. 1), we crossed HSP-12.6::mCherry animals with those expressing a fully-functional N-terminally-tagged myosin heavy chain B, GFP::UNC-54 ^39^, and confirmed that HSP-12.6::mCherry indeed localized to the thick filaments (Fig. 1C). We wondered whether this exclusive localization to thick filaments was due to the higher than endogenous protein levels, even in our low copy-number transgenics. However, recently available endogenously-tagged HSP-12.6::mKate ^40^ protein exhibited identical localization pattern (Fig. 1D). We did not observe any localization to either mitochondria, nucleus, or muscle attachments (both dense bodies and M-lines), or measurable diffuse signal in the cytosol (muscle belly) (Fig. 1E-G and Sup. Fig. 1A,B). Thus, three differently-tagged HSP-12.6 proteins localize exclusively to the thick filaments in the body-wall muscle cells, independent of the developmental stage or feeding status of the animal.

**Figure 1:**
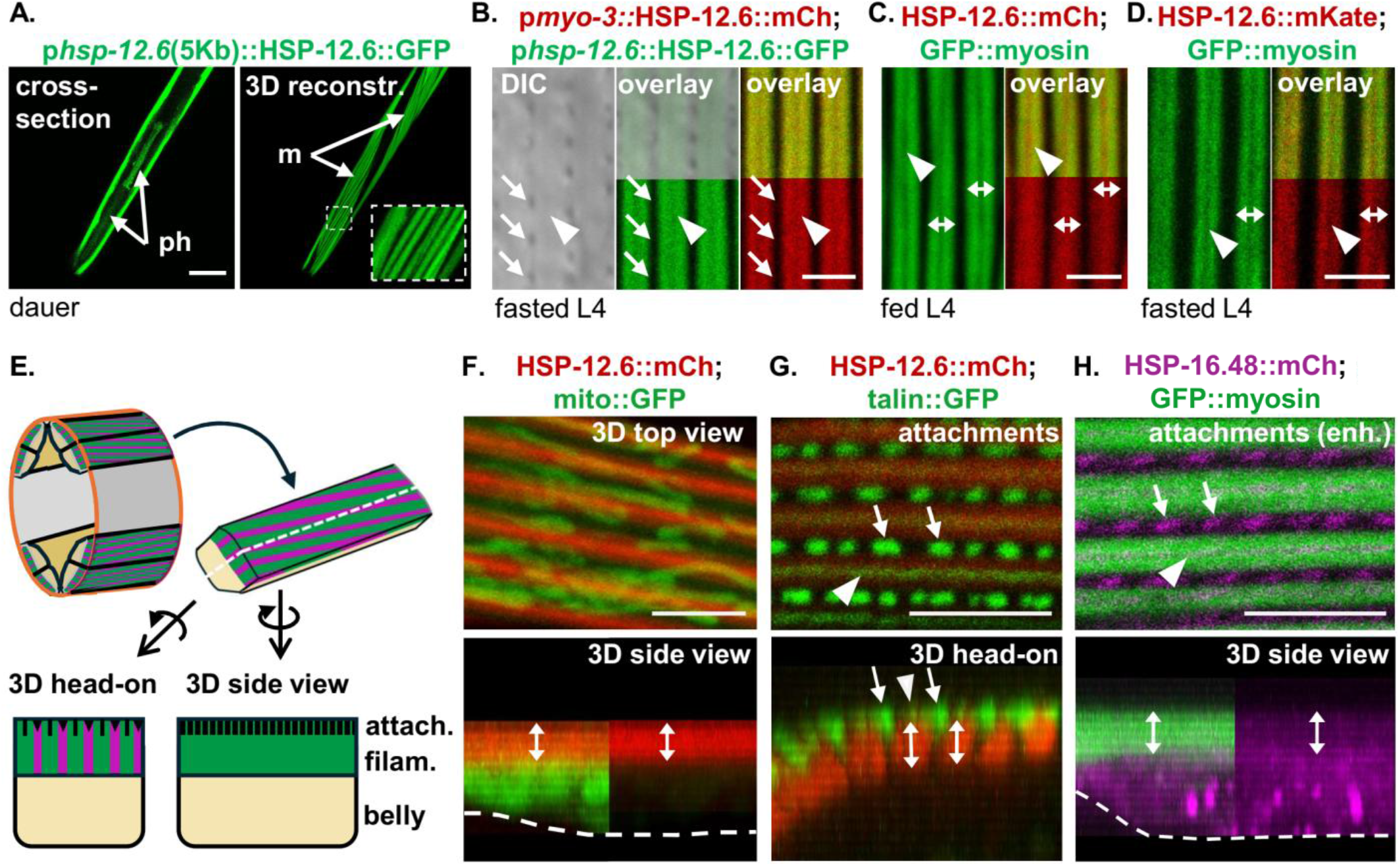
HSP-12.6 has a unique localization pattern to thick filaments in BWM. **A**. Expression of HSP-12.6 in dauers. **Left:** cross-section of the head region. ph: pharynx. **Right:** 3D reconstruction of the same region. m:body-wall muscles; **inset:** striated protein localization. **B-G:** Localization of HSP-12.6::mCherry to thick filaments. Double-headed arrows: filaments; arrows: dense bodies (DBs); arrowheads: M-lines. **B**. GFP-(green) and mCherry-tagged (red) HSP-12.6 share localization to filaments. **C**. HSP-12.6::mCherry localizes to GFP::UNC-54-positive filaments. Myosin: myosin heavy chain B (MHC-B)/UNC-54. **D**. Endogenously-tagged HSP-12.6::mKate localizes as in B. **E**. Schematic of *C. elegans* BWM for orientation of confocal 3D reconstructions in F-H. See also Sup. Fig. 1A. **F**,**G**. HSP-12.6::mCherry (red) does not co-localize with mitochondria (F, mito::GFP, green) or muscle attachments (G, talin::GFP, green). Top panel in G shows projection of attachment planes only. Double-headed arrows show the depth of thick filaments; arrows: DBs; arrowheads: M-lines; punctate line: muscle belly. Note the absence of mCherry signal in the belly. **H**. HSP-16.48::mCherry (magenta) localizes to DBs and M-lines, and diffusely in the muscle belly. Markings as in G. Top panel is brightness enhanced. Scale bars: A: 20µM, B-D: 2.5µM, E-G: 5µM.

This exclusive localization to the thick filaments was unusual, since sHSPs in mammalian muscle cells typically localize to thin filaments and to or near z-disks, through interactions with actin and actin-binding proteins, desmin, and specific areas of titin ^37,38,41,42^. Therefore, we compared HSP-12.6 localization with that of an HSP-16 family protein. HSP-16.48::mCherry protein^27^ exhibited a more typical sarcomeric sHSP pattern, with localization in the attachments plane to both dense bodies and M-lines, and strong diffuse signal throughout all planes of the muscle (Fig. H and Sup. Fig. 1C). Importantly, we detected very little or no enrichment of HSP-16.48::mCherry protein on thick filaments, further highlighting the uniqueness of HSP-12.6 localization. Considering the muscle-restricted expression pattern of HSP-12.6, this raises a question of whether this selective binding reflects its interaction with its physiological client protein(s) in the muscle.

Interestingly, the localization of HSP-12.6 within the thick filament was somewhat variable: in some filaments, it completely overlapped the UNC-54/MHC-B, which is absent from the central-most area of the filament, occupied by the MYO-3/MHC-A protein (Fig. 1 B,D, dark area along the M-line marked with arrowheads; see also scheme in Sup. Fig. 1), while on others it was distributed across entire filament (Fig. 1C). We thus attempted to determine which protein component of the thick filament was responsible for the HSP-12.6 recruitment to the thick filaments. We deleted or downregulated the myosins: UNC-54 and MYO-3, the paramyosin protein UNC-15, and the filament core proteins filagenins. However, HSP-12.6 was still localizing to the remaining thick filaments (Sup. Fig. 1D), suggesting that either a different protein is the client, or that HSP-12.6 recognizes more than a single component of the thick filament.

**Supplemental Figure 1:**
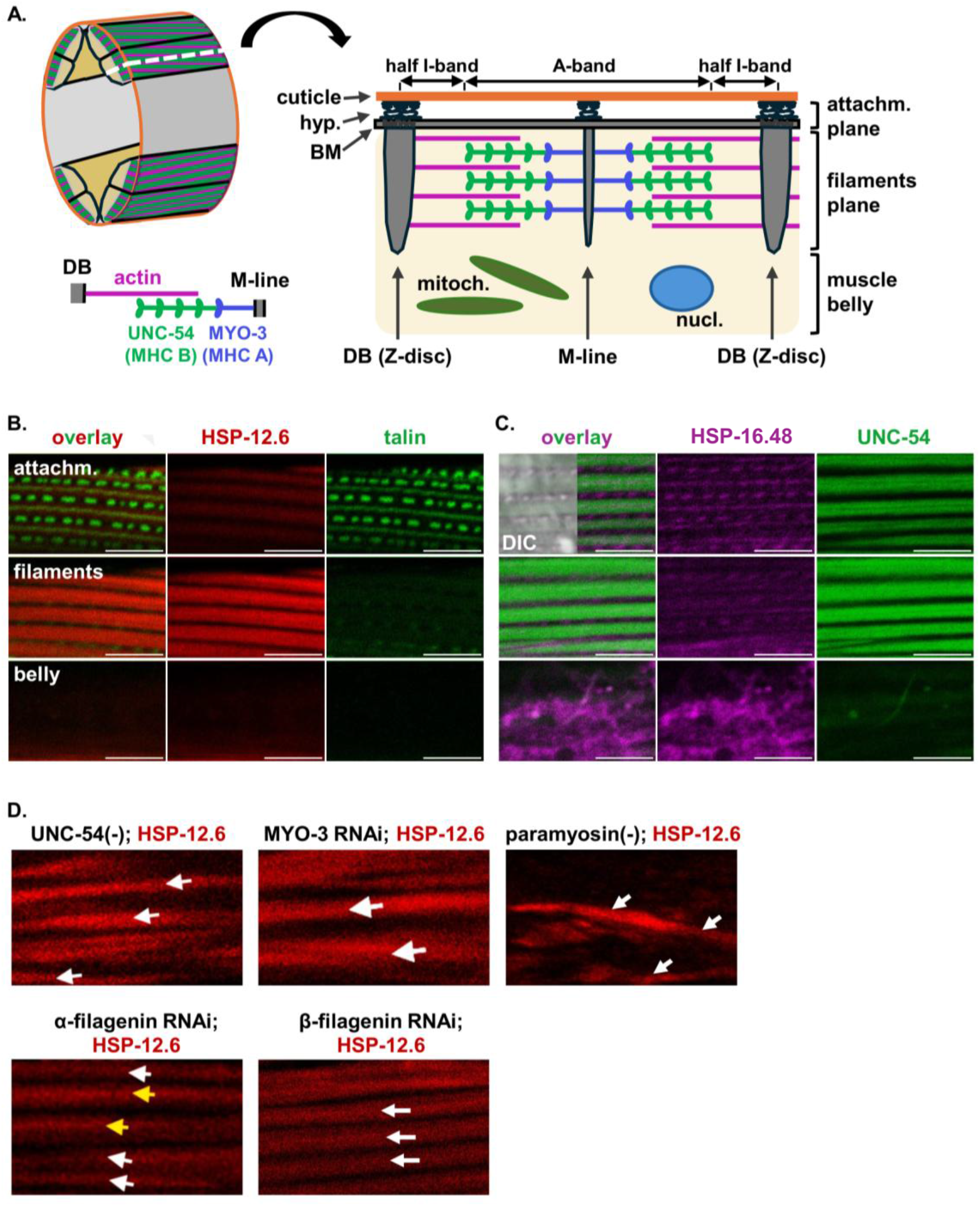
Distinct localization of HSP-12.6 and HSP-16.48 in the muscle cells. **A**. Schematic of *C. elegans* BWM cell and sarcomere organization. Imaging planes are indicated on the right. hyp: hypodermis; BM: basement membrane. **B**. Confocal projections for localization of HSP-12.6 (red) and talin (green) in the attachment plane (top), filaments plane (middle), and muscle belly. **C**. Projections (as in B) of HSP-16.48 (magenta) and MHC-B/UNC-54 (green). B and C are related to Fig. 1G and H; scale bars: 10 µM. **D**. HSP-12.6::mCherry still localizes to myofilaments when indicated thick filament proteins are deleted (-) or downregulated (RNAi).

### HSP-12.6 expression in BWMs selectively rescues severe phenotypes caused by misfolding and mis-assembly of thick filament proteins

Whether localization to the thick filaments in the healthy, unstressed muscles reflects chaperone-client recognition is unclear; moreover, accumulation of sHSPs on sarcomeres or attachments was suggested to represent either a “sink” for excess of sHSPs ^38^, or, in epidermal cells, a reservoir of chaperones poised to relocate to unfolded proteins upon stress ^43^. Therefore, we asked whether the thick filament localization of HSP-12.6 was functional, by testing whether HSP-12.6 can protect against defects caused by misfolding of thick filament proteins. We used the well-characterized temperature sensitive (ts) mutations in paramyosin (UNC-15, a core thick filament protein, homologous to the myosin tail) and myosin (UNC-54), that as previously showed, cause misfolding of these proteins and are sensitive to the state of the muscle proteostasis ^44,45^.

The paramyosin ts mutation (*unc-15(e1402)*) allele causes severe defects in embryo elongation and hatching at the restrictive temperature of 25°C, due to defects in muscle contractions, and decreased motility in surviving animals. We found that introduction of the low-copy HSP-12.6::mCherry into animals expressing the paramyosin(ts) protein strongly suppressed the hatching defect: *paramyosin(ts)*;HSP-12.6::mCherry animals had 5+/5.5% unhatched animals, not significantly different from the wild-type (p>0.999), compared with 45+/-17.6% in *paramyosin(ts)* alone (Fig. 2A). HSP-12.6 also significantly suppressed the movement defect (Fig. 2B).

**Figure 2:**
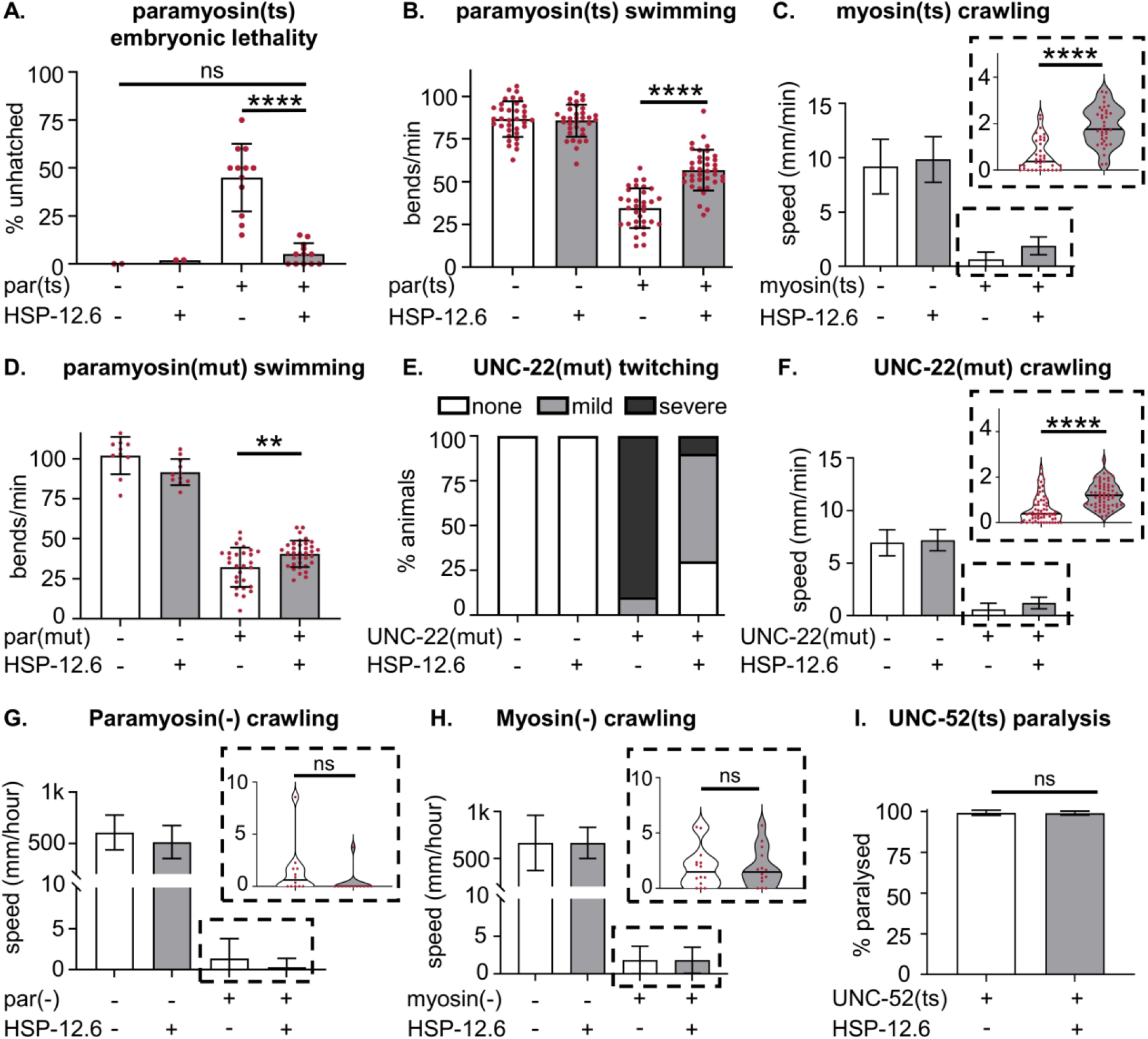
HSP-12.6 expression in BWM rescues severe defects caused by folding and assembly mutations in thick filament proteins. All raw data, n numbers, and p-values are in data supplement; mutant proteins and phenotypes scored are indicated for each panel. **A-C**. HSP-12.6 rescues muscle disfunction caused by folding mutations (ts) in paramyosin (UNC-15) and myosin (UNC-54) proteins. **A**. *unc-15(e1402)*; embryo hatching was scored 24hrs post gastrula stage at 25°C. Data points are individual plates, 20-50 embryos each. **B**. Same strains as in A; animals were exposed to 25°C at L4 stage for 24hrs, and their bending (thrashing) rate when swimming in liquid was measured. **C**. UNC-54(G387R)::GFP L4 animals were exposed to 23°C for 24hrs, placed onto fresh plates in groups of 5, and their crawling speed measured for 1 min. **D-F**. HSP-12.6 rescues disfunction caused by assembly mutation in paramyosin and loss-of-function mutations in thick filament organizing protein twitching/UNC-22 at 20°C. **D**. *unc-15(e1215)*animals; scored as in B. **E**. *unc-22(st136)* animals; scored for 1 min as severe (strong, sustained twitches), mild (occasional twitches), or non-twitching by blinded investigators. **F**. Same strains as in E; scored as in C. **G-H**. HSP-12.6 did not improve crawling in paramyosin(-) (*unc-15(e1214)*) or myosin(-) (*unc-54(e190)*) deletion mutants, scored as in C. **I**. HSP-12.6 did not rescue paralysis in animals expressing mutant UNC-52(ts) basement membrane protein (*unc-52(e669su250)* allele), day1 adults grown at 20°C. Data points in B-D and F-H are individual animals.

The activity of HSP-12.6 did not depend on stressful temperature, since it also rescued the milder hatching defect of *paramyosin(ts)* animals at the normal temperature of 20°C, as well as its developmental delay in animals grown at an intermediate temperature of 22°C (Sup. Fig. 2A-B). We noticed, however, that animals with HSP-12.6 alone grew somewhat slower than non-transgenic animals (N2). We then scored them at the time when N2 animals have just became reproductive (97+/-3.1%, at 65-66 hrs at 20°C), and found that only 1+/-1% of HSP-12.6 animals were reproductive, with the rest being either young adults (82+/-2%, up to 3 hrs delay) or in the fourth larval stage (17+/-2%, L4, up to12 hrs delay) (Sup. Fig. 2C). Thus, while HSP-12.6::mCherry was able to *decrease* the developmental delay in animals with the folding paramyosin mutation, it *caused* a mild delay in otherwise wild-type animals, suggesting that its expression in the absence of a ‘needy’ client is in fact detrimental.

We confirmed the ability of HSP-12.6 to rescue the effects of myofilament protein misfolding using a ts mutant myosin protein, UNC-54. The G387R mutation, identical to that of the destabilized *unc-54(e1301)* allele ^45^, is knocked-in to the endogenously-tagged UNC-54::GFP ^46^; we chose this strain to allow for subsequent direct imaging of mutant myosin aggregates (in Fig. 4). UNC-54(ts)::GFP animals have low crawling speed at an intermediate temperature, 23°C, and HSP-12.6::mCherry was able to partially suppress this defect (Fig. 2C). Interestingly, while the rescue of the mean crawling speed was statistically significant but of a rather small magnitude, we observed a complete rescue of paralysis (inset in Fig. 2C).

We then asked if HSP-12.6 could rescue the defects caused by mutations affecting thick filament assembly or organization, using an assembly mutation in paramyosin (UNC-15, *unc-15(e1215)* allele), and a loss of function mutation in myofilament organization protein, twitchin (UNC-22, *unc-22(st136)*). HSP-12.6::mCherry moderately rescued the swimming defect in paramyosin assembly mutants (Fig. 2D), and improved the crawling speed in *unc-22* mutants (Fig 2F). Strikingly, like with the myosin ts mutation, expression of HSP-12.6::mCherry effectively suppressed the most severe phenotypes in *unc-22* mutants: it eliminated paralysis (inset in Fig. 2F), by strongly decreasing the severe twitching phenotype (p<0.0001, Fig. 2E).

**Supplemental Figure 2:**
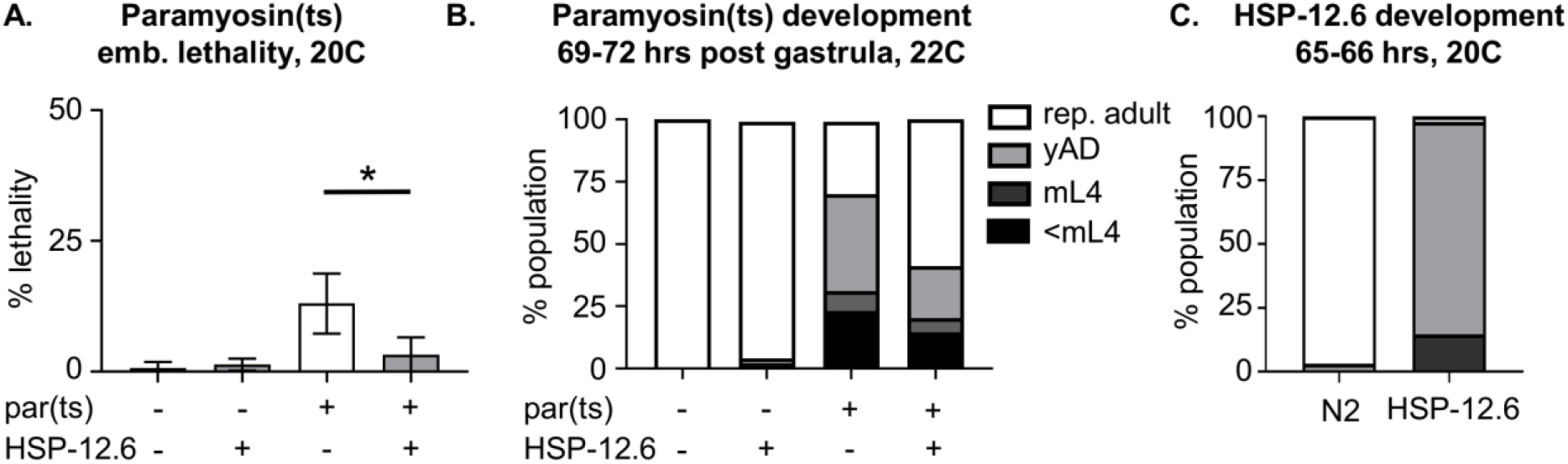
**A**. HSP-12.6 rescues hatching defect at normal growth temperature, 20°C. Scored as in Fig. 1A. **B**. HSP-12.6 improves development defect in paramyosin(ts) animals at 22°C; scored at 72 hours post gastrula. **C**. HSP-12.6 animals have mild developmental delay when expressed without a ‘needy’ client. Scored at ∼65.5 hrs at 20°C.

The observed rescue of defects caused by misfolded or misassembled thick filament proteins could be a result of a general increase in muscle health or functionality, rather than of a client-dependent role of HSP-12.6. Thus, we asked whether it can rescue defects in animals with a complete loss of paramyosin or myosin proteins, and in those with misfolded non-myofilament proteins. HSP-12.6 expression failed to rescue the severe movement defects of paramyosin (*unc-15(e1214)*) or myosin/UNC-54 (*unc-54(e190)*) deletion alleles. Moreover, in contrast to its rescue of paralysis in folding/assembly mutants, its expression actually increased the number of completely paralyzed paramyosin deletion animals (inset in Fig. 2G). Similarly, HSP-12.6 did not decrease the percent of paralyzed animals with the folding (ts) mutation in perlecan/UNC-52 protein (*unc-52(e669su250)*) (Fig. 2I) grown at 20°C, or the 100% resistance to the nicotinic acetylcholine receptor agonist levamisole in animals with a folding mutation in the muscle-expressed receptor subunit (*unc-63(x26)*), at the restrictive 25°C.

Therefore, HSP-12.6 possesses a client-dependent protective activity against proteostatic mutations in thick filament proteins, and efficiently rescues their severe phenotypes.

### HSP-12.6 cannot rescue phenotypes associated with thin filament mutations

We were curious whether HSP-12.6 selective localization to the thick filaments reflected its selective function, or whether it was able to support all myofilaments. Indeed, since the muscle organization depends on the ordered interaction between both types of filaments, disruption of one type may indirectly affect the organization of the other. We used mutations in the thin filament assembly protein UNC-78 (orthologous to the human WD repeat domain 1, *unc-78(e1217)* allele), and a calponin-related UNC-87 (*unc-87(e1459)*), which maintains organization of thin filaments ^47,48^. We found that HSP-12.6 failed to rescue movement in either of these mutants (Fig. 3), supporting the idea that the thick-filament exclusive localization of HSP-12.6 reflects its selective activity.

**Figure 3:**
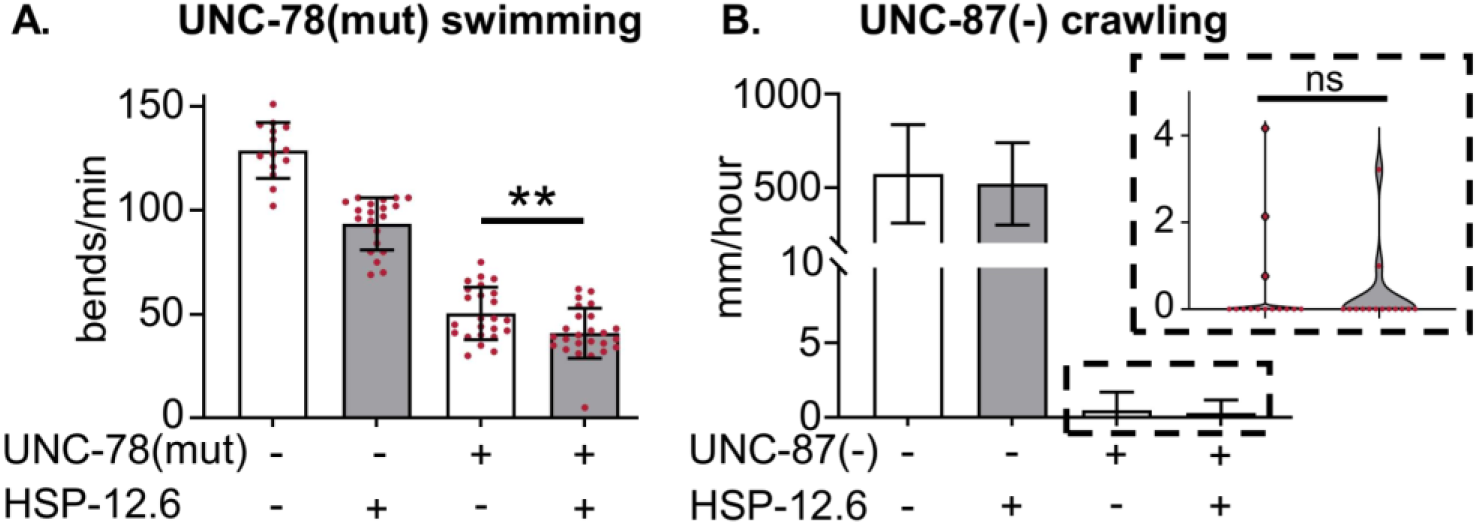
HSP-12.6 cannot rescue muscle defects associated with thin filament mutations. **A**. HSP-12.6 significantly increases swimming defect in males with a mutant thin filament assembly protein UNC-78 (*unc-78(e1217)* allele). Scored as in Fig 2B. **B**. HSP-12.6 does not rescue crawling deficit in animals with a mutant thin filament organizing protein UNC-87 (*unc-87(e1459)* allele). Scored as in Fig 2C.

**Figure 4:**
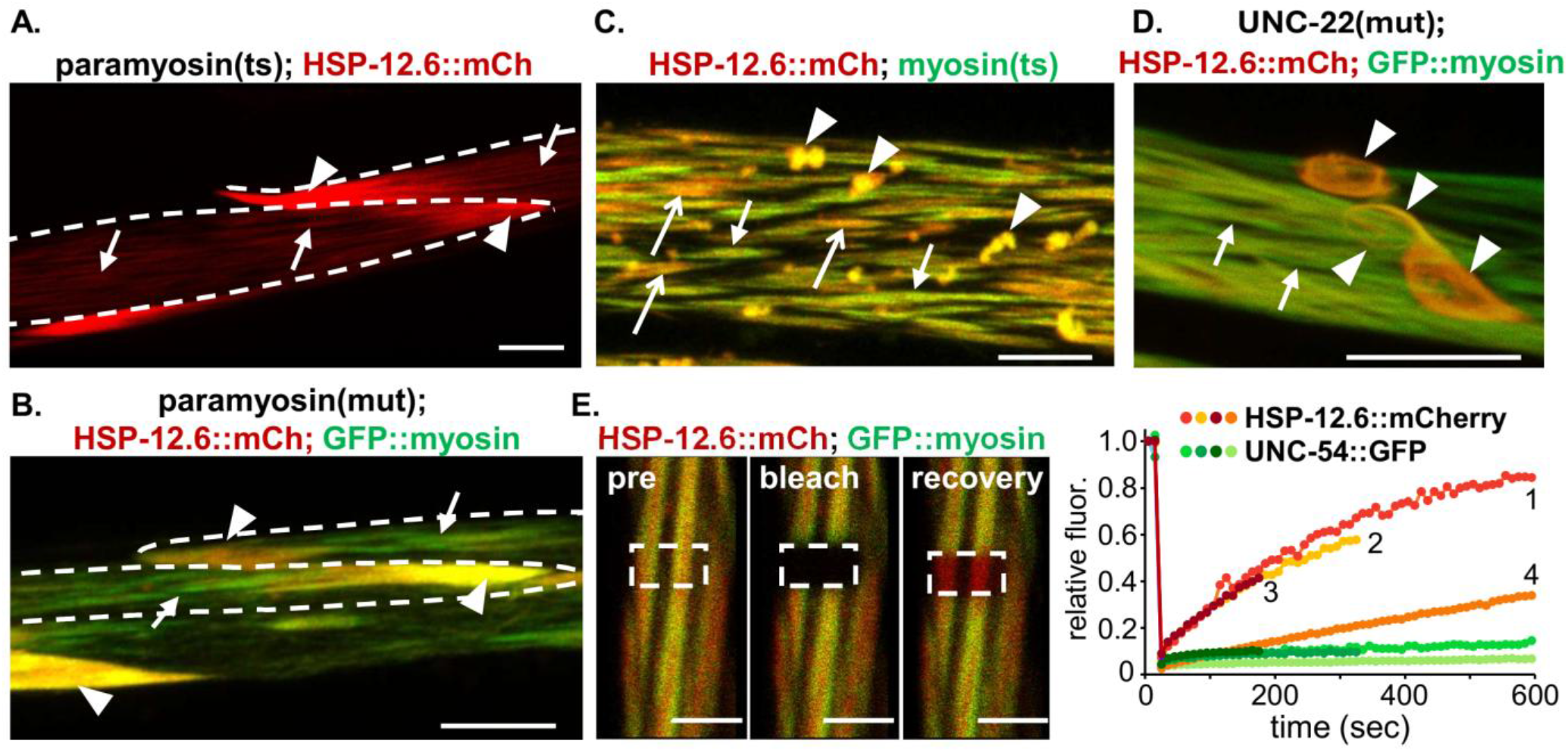
HSP-12.6 localizes to thick filament proteins with chaperone-like properties. In panels A-D, arrowheads mark aggregates; arrows and open arrows mark remaining filaments with weak or strong HSP-12.6::mCherry binding, respectively. Proteins expressed are indicated on top. **A-B**. HSP-12.6::mCherry (red) re-localizes to needle-like aggregates at the tips of muscle cells (punctate outlines) in animals expressing folding (ts) or assembly mutant paramyosin. **A**. L4 larvae were shifted to 25°C for 20 hrs and live imaged. Remaining thick filaments are only weakly positive for HSP-12.6::mCherry. **B**. GFP::myosin (GFP::UNC-54, green) is present in both remaining filaments and aggregates, while HSP-12.6::mCherry (red) is predominantly in aggregates. **C**. HSP-12.6::mCherry (red) similarly re-localizes to globular aggregates of myosin(ts) (UNC-54(ts)::GFP, green); myosin(ts) is also present in the remaining filaments with weak or strong HSP-12.6::mCherry binding. L4 animals exposed to 25°C for 16 hrs. **D**. Mutation in thick filament organizing protein UNC-22 induces abnormal coiled filaments and filamentous aggregates. HSP-12.6::mCherry relocalizes to both. **E**. HSP-12.6::mCherry fluorescence (red) recovers after photobleaching, while GFP::UNC-54 fluorescence (green) does not. Left panels: indicated region of interest (ROI) before and immediately after simultaneous GFP/mCherry photobleaching, and after recovery. HSP-12.6 (red) is visible in the ROI after recovery. Right: Bleach/recovery graphs for four cells; imaging stopped when animals moved (curves 2 and 3); curve 4 is of a head muscle.

Since the movement defect in *unc-78* mutants has a later onset, it forced us to use male animals to avoid the confounding effects of egg retention in hermaphrodites; interestingly, we observed a tendency for decreased motility in both wild type and mutant male animals expressing HSP-12.6, with the mutant animals reaching statistical significance (Fig. 3B). While the exact reason for this is presently unknown, it is possible that hyperactive movement, characteristic of males, makes their muscle environment more sensitive.

### HSP-12.6 localizes to thick filament proteins with chaperone-like properties

Our data so far is suggestive of a thick-filament – selective chaperone function for HSP-12.6. If it indeed binds thick filaments as a chaperone, we would expect preferential binding to misfolded clients over healthy proteins, as well as their interactions being transient, with slow kinetics. To test for preferential binding, we examined HSP-12.6::mCherry localization in animals carrying the folding/assembly mutations in thick filament proteins.

Many mutations in paramyosin are known to cause accumulation of paracrystalline-like aggregates, often referred to as ‘needles’, in the distal corners of muscle cells ^49^; in addition to paramyosin, these aggregates contain other thick-filament-associated proteins ^50-53^. We found a striking redistribution of HSP-12.6 from remaining filaments to these aggregates in muscles with both the folding (ts) and assembly mutants of paramyosin/UNC-15 protein (Fig. 4A,B). In addition, when GFP::UNC-54 myosin was included in animals with the assembly mutant of paramyosin, it was readily detected both in both the needles and in the remaining disorganized thick filaments, while HSP-12.6::mCherry was predominantly detected in the needles (compare the mostly-green filaments (arrows) and intensely yellow needle-like aggregates (arrowheads) in Fig. 4B).

Similarly, in animals carrying the folding mutation in the endogenously-tagged UNC-54(ts)::GFP, HSP-12.6::mCherry was localizing strongly to the GFP-positive globular aggregates (Fig. 4C, arrowheads) at restrictive temperature. Interestingly, in these animals, some of the remaining thick filaments were also intensely positive for HSP-12.6 (open arrows), suggesting that the presence of misfolded myosin did increase its binding. Finally, we examined the muscle in animals carrying *unc-22* myofilament organization mutation, along with GFP-UNC-54 to visualize filament (Fig. 4D); we found that muscles in these animals, in addition to the disorganized filaments (arrows), contained long and abnormally shaped or coiled UNC-54/myosin-positive filaments and aggregates (arrowheads). Both the aggregates and the abnormal filaments were strongly positive for HSP-12.6::mCherry. Thus, HSP-12.6 preferentially binds aggregates and assemblies containing non-native thick filament proteins.

Because UNC-54 myosin (MHC-B) was recruited into the needle-like aggregates caused by the paramyosin mutation, we asked if this was also true for the MHC-A protein, MYO-3. We found that while, as expected, the needles were intensely positive for HSP-12.6::mCherry, GFP::MYO-3 protein was excluded from the needles (Supl. Fig. 3), uncovering a remarkable difference in the interactions of the two myosins with non-native proteins in these animals. Together with our RNAi findings in Sup. Fig. 1D, this shows that MYO-3 is not the client that recruits HSP-12.6 to either healthy filaments or aggregates.

**Supplemental Figure 3:**
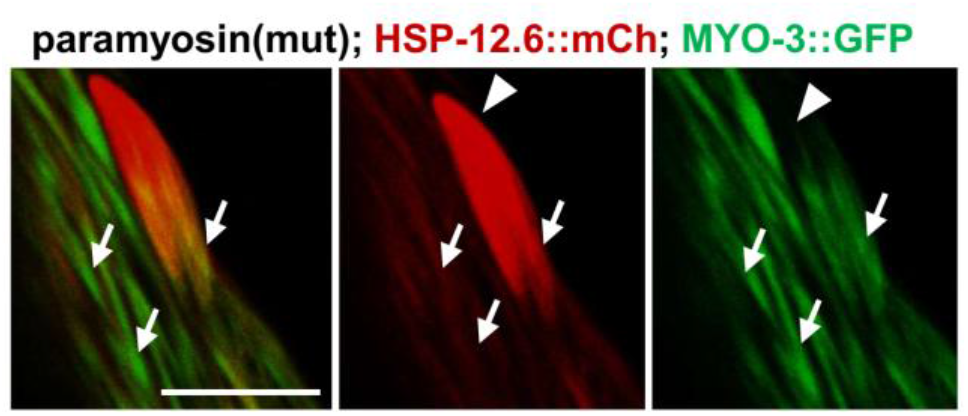
Myosin heavy chain A (MYO-3::GFP, green) is absent from HSP-12.6::mCherry-positive aggregates (arrowhead) and remains within the disrupted filaments (arrows) in paramyosin assembly mutants.

The second criterion for the chaperone-client interaction is their transient binding. To test this, we performed FRAP analysis in the muscles of HSP-12.6::mCherry;GFP::UNC-54 animals. When a section of the sarcomeres was photobleached, the fluorescence of HSP-12.6::mCherry showed consistent recovery (red to orange curves 1-3, Fig. 4E), indicating that its interaction with thick filaments was dynamic. Conversely, the fluorescence of GFP::UNC-54 myosin in the same bleached region never recovered (over 60 min), consistent with this protein being a structural component of the filament (green curves). The recovery of HSP-12.6 was surprisingly slow, precluding us from measuring the completeness of recovery due to movement of the animals; however, we were able to live image one BWM cell without large movements for 60 min, measuring recovery to above 80% relative to pre-bleach levels (Fig. 4E, curve 1). The slow kinetics of recovery are likely explained by the absence of the readily available soluble pool for exchange, as judged by the exclusive localization of HSP-12.6::mCherry to filaments and the absence of diffuse signal. In this case, the only source of unbleached protein for the recovery of fluorescence will be proteins that dissociated from unbleached areas of thick filaments in the same cell. In agreement, when we performed FRAP in one of the head muscles, which have lower expression of the HSP-12.6::mCherry transgene, the recovery was even slower (Fig. 4E, curve 4).

The strong preferential binding of HSP-12.6 to non-native assemblies, whether they were globular aggregates, organized needle-like aggregates, or abnormal filaments, prompted us to ask whether it would similarly bind to aggregates that were not generated by the thick filament proteins. Muscle cells of *unc-78* mutant animals are known to contain aggregates that are positive for actin and some actin-binding proteins, and are morphologically similar to the paramyosin-containing needles ^47^. We confirmed the presence of such aggregates in our *unc-78* mutants, using actin-binding protein CLIK-1::GFP, and found that HSP-12.6::mCherry is excluded from these aggregates (Fig. 5A). Although this is in agreement with our rescue experiments, and with exclusion of HSP-12.6 from thin filaments in non-stressed muscles, such dramatic selectivity in binding to non-native proteins and/or their aggregates is still surprising when considering sHSP chaperones. We thus asked a reciprocal question, and found that HSP-16.48::mCherry did not bind the UNC-54/myosin-positive needle-like aggregates in paramyosin assembly mutants (Sup. Fig. 4). Given that HSP-16.48 is a broad-specificity sHSP, this may indicate that the needle-like aggregates are sites of inappropriate assembly of thick filament proteins, rather than aggregates of misfolded proteins, and that HSP-12.6 preferentially interacts with both the misfolded and mis-assembled thick filament proteins.

**Figure 5:**
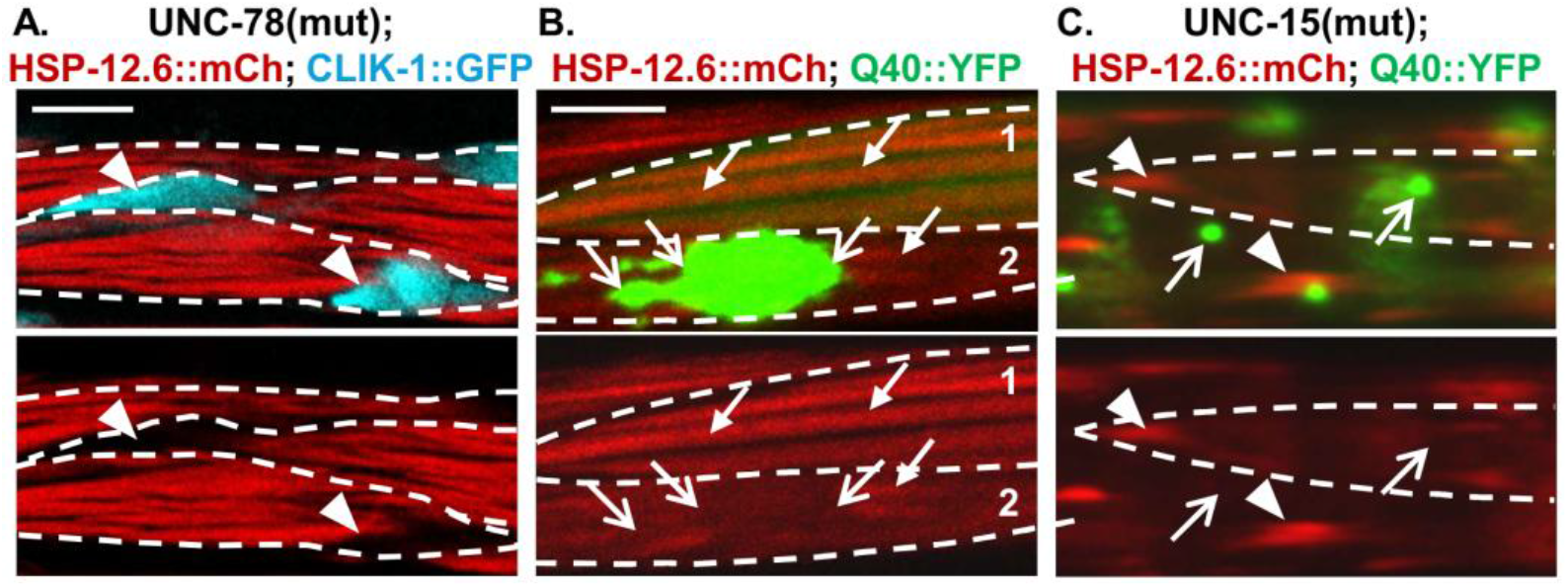
HSP-12.6 does not recognize aggregates of thin filament proteins, and selectively binds thick filaments or their aggregates over polyQ40. In all panels, arrowheads: needle-like aggregates; arrows: thick filaments; open arrows: polyQ40 aggregates. Proteins expressed are indicated on top. **A**. HSP-12.6::mCherry (red) is excluded from actin-containing needle-like aggregates induced by the mutant thin filament assembly protein UNC-78. Cyan: actin-binding protein CLIK-1::GFP. **B**. HSP-12.6::mCherry continues binding thick filaments in cells with either soluble (cell 1) or aggregated (cell 2) polyQ40::YFP and does not re-localize to polyQ40 aggregates. Early aggregates in L4 animals. **C**. HSP-12.6::mCherry localizes to the paramyosin needles over polyQ40 aggregates even when challenged with both the paramyosin and polyQ40 aggregates in the same cell (punctate outline).

Finally, we asked whether HSP-12.6 would retain its selectivity if challenged with a disease-related aggregation-prone protein. We used a polyglutamine expansion Q40::YFP, since we previously found it to disrupt the proteostasis in the muscle and to expose the ts phenotypes of both paramyosin and myosin mutants at permissive temperatures ^45^. HSP-12.6::mCherry did not localize to the polyQ40 aggregates (Fig. 5B, arrowheads), remaining instead bound to the myofilament in cells with either soluble or aggregated Q40 (arrows). Moreover, it retained its remarkable selectivity when presented with both the polyQ40 and the thick filament aggregates in the same cell, using *unc-15(e1215);*HSP-12.6::mCherry;Q40::YFP animals: HSP-12.6::mCherry was strongly localized to the needle-like aggregates (Fig. 5C, arrows), while exhibiting no localization to the polyQ40::YFP aggregates (arrowheads).

**Supplemental Figure 4:**
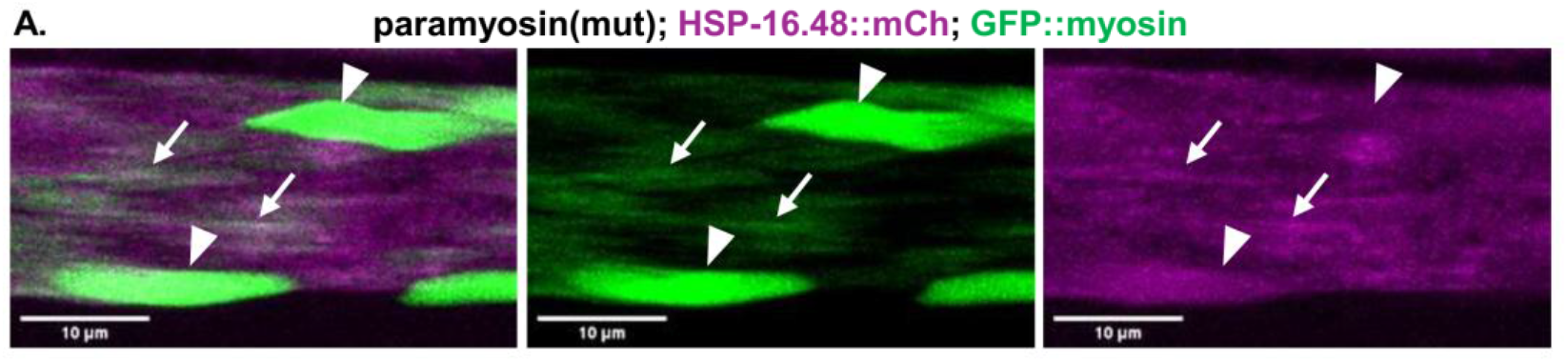
HSP-16.48 does not localize to the needle-like aggregates in paramyosin assembly mutants. HSP-16.48 remains diffuse, with some filament localization (arrows), and is absent from the thick filament needle-like aggregates (arrowheads). Compare to HSP-12.6 in Fig. 4B.

## Discussion

Our findings resolve a long-standing conundrum – the combination of strong genetic evidence for important *in vivo* function(s) for the small heat-shock protein HSP-12.6 with its complete lack of activity in chaperone assays. We find that activity of HSP-12.6 *in vivo* is directed towards a highly restricted set of clients (or a client) in the tissue where its expression is controlled by development or by physiological conditions – the body-wall muscles. Indeed, HSP-12.6 consistently exhibited a myoprotective function, but only when the muscles were challenged with mutations causing misfolding or mis-assembly/aggregation of thick filament proteins, or disruption of thick filament organization. In contrast, it showed no activity not only towards the generic misfolded (ts) proteins, but also towards mis-assembled/aggregated thin filament proteins and disrupted thin filaments, confirming that HSP-12.6 is truly client(s) selective. As has been proposed for the narrowly-expressed mammalian specialist sHSPs with no activity in chaperone assays, this selectivity likely explains the lack of HSP-12.6 chaperone function towards model unfolded substrates, or when expressed in non-cognate cells. It would be informative thus to examine the other three members of the HSP-12 family, of which two – HSP-12.2 and HSP-12.3 were tested *in vitro* and found to be similarly inactive ^25^, for their expression patterns and binding/chaperone preferences.

The combination of fluorescent tagging and live imaging allowed us to detect HSP-12.6 binding preferences in both healthy unstressed animals and in animals with non-native client proteins. HSP-12.6 interacted exclusively with the native thick myofilaments, with binding properties consistent with the client-chaperone interaction, and relocalized to their non-native assemblies or aggregates in mutant animals. When challenged with deliberately introduced polyglutamine expansions, which as we have previously shown ^45^, disrupt the muscle proteostasis, HSP-12.6 selectively bound to thick filament protein aggregates over polyQ40 aggregates, even when both were present in the same cell. This indicates continued exclusive binding despite the exceptionally high proteotoxic stress, and my explain the lack of detection of any HSP-12 family proteins in the aging-induced protein aggregates, which contain several other *C. elegans* sHSPs ^28,29^.

Remarkably, HSP-12.6 localized to non-native assemblies of thick filament proteins regardless of their nature – from globular aggregates triggered by misfolded myosin, to the ordered needle-like aggregates in paramyosin mutants, to abnormal filaments in *unc-22* mutants. This raises the question of what this chaperone actually recognizes. One possibility is that it recognizes all conformations of its client(s), including the native conformation in healthy filaments. This is difficult to reconcile with its stark client selectivity, unless the client retains the native conformation of the site recognized by HSP-12.6 even within aggregates.

Second, since our attempt to pinpoint the protein that recruits HSP-12.6 showed that none of the tested thick filament structural proteins alone are responsible, it is possible that HSP-12.6 binds filament-interacting protein(s), rather than filaments themselves. This would be similar to the mammalian HSPB1 and HSPB5 binding to titin or desmin both in aggregates and in Z-disks and I-bands of unstressed muscles ^37,38,42^, the latter likely by binding to locally unfolding (due to mechanical strain) domains of these proteins ^54,55^. An interesting candidate with similar properties but in the thick filaments would be MYBPC3 ^56^, however, its homologue has not been identified in *C. elegans*.

Finally, it is also possible that the HSP-12.6 – positive aggregates contain some amount of assembled normal thick filaments, or perhaps their fragments. This is quite likely for the needle-like aggregates, since similar, if smaller, paramyosin needles were found to contain ordered filamentous structures ^49^. These aggregates being sites of mis-assembly is consistent with the exclusion of the myosin MYO-3/MHC-A from the needles that are positive for both paramyosin^49^ and the myosin UNC-54/MHC-B, since these two myosins assemble in the distinct domains of the thick filament (see scheme in Sup. Fig. 1). This is also consistent with the failure of the broad-selectivity sHSP, HSP-16.48, to bind the thick filament needles, suggesting that they may not include misfolded proteins. However, it is unclear if the HSP-12.6 – positive globular aggregates induced by the myosin(ts) misfolding contain such filamentous fragments. Future work will be needed to identify the client of HSP-12.6 in the thick filament, which will then allow determination of the consequences of their interaction to the client.

In several assays, we observed a detrimental effect of HSP-12.6 expression in non-mutant animals on the muscle function, including slowing of development and movement; similar detrimental effects have been reported for overexpression of chaperones, for example of HSP70 in *Drosophila* ^57^. Moreover, in the presence of proteostatic challenge to the non-client thin filaments, HSP-12.6 expression potentiated the toxic phenotype. Although not overexpressed, our HSP-12.6::mCherry transgenic protein is expressed in the muscles at all developmental stages, while endogenous expression is seen in the first larval stage (L1) and in dauers ^32^, and in fasted or starved animals ^35,36^. This may suggest that its function is specifically needed at these stages/conditions. L1 larvae switch from restricted movement in the embryo to crawling, as well as growth of the muscle cells and addition of sarcomeres ^58,59^, although sarcomere addition continues to happen in the rest of the larval stages ^59^. Dauer muscles are remodeled to have larger sarcomeres; in addition, while undisturbed dauers exhibit little movement, they are capable of bursts of intense movement when stimulated ^60^. It is possible then that HSP-12.6 is specifically needed to support the myofilaments undergoing remodeling, or during increased movement, although we observed a stronger detrimental effect in the more movement-active males than in hermaphrodites. An important consideration of these findings is that proposed approaches to activate sHSPs as treatment possibilities in protein aggregation myopathies ^61^ may need to be targeted to the affected cells only, and/or stratified by the nature of the damaged proteins and the sHSP being targeted.

The high proportion of mammalian sHSPs being expressed in the different muscle cells, as well as their involvement in myopathies, suggests that the muscles have generally high needs for these chaperones. Indeed, many sarcomeric proteins are continuously subjected to mechanical strain, while conditions like energy demands and calcium fluctuations create an environment that challenges proteostasis ^62,63^. It may also suggest that at least for some sHSPs, muscle-enrichment reflects tissue-distribution of their specific clients. For example, Filamin C, the client of cardio-vascular HSPB7, is preferentially expressed in cardiac and skeletal muscles, where it functions by organizing actin-containing thin filaments ^20^. The muscle-specific expression pattern of HSP-12.6 expression, and its ability to decrease or even prevent the muscle dysfunction in thick filament mutant animals, places it into such myoprotective chaperone class. Its selectivity to the thick filaments, however, distinguishes it from most of the other sHSPs; while HSPB1 and HSPB5 have been shown to interact with myosin, this was with deliberately unfolded myosin ^64,65^. Interestingly, in myopathies, sHSPs are often detected in aggregates containing thin filament/Z-disc and intermediate filament proteins, but have not been found within myosin-positive aggregates ^66-70^; we have similarly detected no binding of HSP-16.48 to the thick filament aggregates. Myosin-storage myopathies are the least mechanistically understood group ^71-73^, and thus it would be important to determine if any of the specialist human sHSPs share the thick-filament selectivity of HSP-12.6, and whether its selective myoprotective properties can be leveraged against such myopathies.

## Materials and Methods

### Nematode strains and genetics

Standard methods were used for worm culture and genetics ^74^. Animals were kept at constant 20°C temperature throughout, unless specifically indicated. Temperature sensitive animals were grown and maintained at 15°C and shifted to indicated temperatures for experiments. D1A animals were obtained by picking L4 and scoring 24 hours later,

The following stains were obtained from *Caenorhabditis* Genetics Center (CGC): AH3437(*tln-1(zh117[gfp::tln-1]*)I), CB190(*unc-54(e190)* I), CB1215(*unc-15(e1215)* I), CB1214(*unc-15(e1214)* I), CB1217(*unc-78(e1217)* X), CB1402(*unc-15(e1402)* I), CB1459(*unc-87(e1459)* I), HE250(*unc-52(e669su250)* II), NIS1013(*kytEx1013*[p*hsp-12*.*6(5kb)*::*hsp-12*.*6*::GFP + *rol-6(su1006)*]), NL3643(*unc-22(st136)* IV), ON352(*clik-1(kt1[clik-1::gfp])* V), PD2882(*unc-54(G387R::gfp::TAA::NSUTR)* I), PXH1364(*hsp-12*.*6(syb136[hsp-12*.*6::mKate2])*IV), PD4251(*ccIs4251[(pSAK2) myo-3p::GFP::LacZ::NLS + (pSAK4) myo-3p::mitochondrial GFP + dpy-20(+)] I)*.

The AS408[p*unc-54*::GFP::UNC-54] and AM134(*rmIs126*[p*unc-54*::Q0::YFP]) strains were kindly provided by the Morimoto lab, the CNK401(*hsp-16*.*48/49(prp60[hsp-16*.*48/49::mCherry])*) by the Nussbaum Lab, and the KAR007[*drxIs1*(p*myo-3*::*hsp-12*.*6*::mCherry)] by the Rodriguez lab.

The *drxIs1* transgene was made by amplifying a 500bp fragment containing coding region of *hsp-12*.*6* from N2 genomic DNA. *myo-3* promoter for expression in the body-wall muscle cells, worm mCherry, and *unc-54* 3’UTR were amplified from pCFJ104 (Addgene #19328). Fragments were assembled into pMCS5 (MoBiTec, Germany). *drxIs1* was integrated by bombardment as 20ng/uL plasmid DNA and 80ng/uL digested worm genomic DNA by the Rodriguez Lab (UT Health San Antonio).

### Rescue assays

All temperature sensitive (ts) animals were grown and maintained at 15°C. Only animals that had not been starved/crowded or contaminated for at least three generations were used. For restrictive temperatures, animals were shifted from 15°C to 25°C for the indicated amount of time. For intermediate temperatures, pilot experiments were run to determine temperature that provide intermediate phenotypes, mainly between 20-23°C.

### Development Assays

Animals were synchronized by picking gastrula stage embryos and scored after indicated number of hours based on physical characteristics, *i*.*e*. vulva formation, as 4 groups: reproductive adult (Rep Ad), young adult (YA), mature L4 (mL4), or younger than mature L4 (<mL4). Embryo hatching was scored at 24 hrs post gastrula at 25°C.

### Motility Assays

For swimming, synchronized animals at indicated age were placed in a drop of M9 buffer and acclimated for 1 min; the completed body bends were counted using the wrmtrck plugin in ImageJ. At least 10 animals per trial.

For crawling, animals were placed on fresh OP50 bacteria lawn in groups of 5 and allowed to crawl for either 1 min or 1 hour, depending on severity of phenotype. The tracks were imaged and measured using ImageJ; at least 10 animals per data point. For 1min assays, nematodes were stimulated with blue light. For 1 hr assays, the nematodes were unstimulated; wild type animals were measured for 1 minute and the result multiplied by 60. All measurements were taken at the same magnification.

For twitching, D1A *unc-22(st136)* were scored as: 0-no twitching, 1-mild twitching, 2-severe twitching. Each of the three trials was scored blindly by separate individuals.

### Microscopy

For confocal images, animals were immobilized on 2% agar pads with 20 mM Sodium Azide and imaged with Zeiss LSM700 microscope at Cell Imaging Center, Drexel University. Z-stacks were acquired at 0.4 µm intervals as 12-bit images, using 63x 1.4NA objective, and analyzed with ImageJ.

Stereo images and movies were taken using Leica M205FA stereo microscope with Orca R2 digital camera (Hamamatsu). The magnification was kept constant within experiments.

## Statistical analyses

All ANOVA and *t*-test analyses were performed using Prism software (GraphPad, USA). ANOVA was followed by multiple comparison Bonferroni’s post-test’ resulting significance levels are indicated in the figures. α=0.05 was used for all analyses. When means are shown, data are presented as mean+/-st. dev. All data points and exact p-values are included in the data supplement.

## Acknowledgements

We are grateful to Drs. Morimoto, Ruvinsky, and Bott (Northwestern University), Nussbaum (LMU Munich), and Ben-Zvi (Ben-Gurion University) for helpful discussions, and Drs. Morimoto, Nussbaum, and Rodriguez (UTHSA) for sharing strains. Some strains were provided by the CGC, which is funded by NIH Office of Research Infrastructure Programs (P40 OD010440).

Confocal experiments were conducted at Drexel University’s Cell Imaging Center, RRID:SCR_022689.

## Funding

This work was supported by a R36 AG045411 grant to Jasmine Alexander-Floyd, and the Louis and Bessie Stein Family Fellowship to Tali Gidalevitz.

## Notes

### Competing Interest Statement

The authors have declared no competing interest.

